# Activation of the Aryl Hydrocarbon Receptor in Endothelial Cells Impairs Ischemic Angiogenesis in Chronic Kidney Disease

**DOI:** 10.1101/2023.07.24.550410

**Authors:** Victoria R. Palzkill, Jianna Tan, Qingping Yang, Juliana Morcos, Orlando Laitano, Terence E. Ryan

**Author notes:** Correspondence: Terence E. Ryan, PhD: 1864 Stadium Rd, Gainesville, FL, 32611.; Twitter: @TerenceRyan_PhD. **Conflict of Interest Statement**: The authors have declared that no conflict of interest exists.

## Abstract

**Rationale:** Chronic kidney disease (CKD) is a strong risk factor for peripheral artery disease (PAD) that is associated with worsened clinical outcomes. CKD leads to accumulation of tryptophan metabolites that associate with adverse limb events in PAD and are ligands of the aryl hydrocarbon receptor (AHR) which may regulate ischemic angiogenesis.

**Objectives:** To test if endothelial cell-specific deletion of the AHR (AHR^ecKO^) alters ischemic angiogenesis and limb function in mice with CKD subjected to femoral artery ligation.

**Findings:** Male AHR^ecKO^ mice with CKD displayed better limb perfusion recovery and enhanced ischemic angiogenesis compared to wildtype mice with CKD. However, the improved limb perfusion did not result in better muscle performance. In contrast to male mice, deletion of the AHR in female mice with CKD had no impact on perfusion recovery or angiogenesis. Using primary endothelial cells from male and female mice, treatment with indoxyl sulfate uncovered sex-dependent differences in AHR activating potential and RNA sequencing revealed wide ranging sex-differences in angiogenic signaling pathways.

**Conclusion:** Endothelium-specific deletion of the AHR improved ischemic angiogenesis in male, but not female, mice with CKD. There are sex-dependent differences in *Ahr* activating potential within endothelial cells that are independent of sex hormones.

## INTRODUCTION

Peripheral arterial disease (PAD) is a chronic disease primarily affecting the lower extremities which is caused by atherosclerotic narrowing or occlusion of blood vessels. An estimated 8-12 million adults in the United States and ∼200 million globally that have PAD (1–3). The obstruction of blood flow in the lower extremities can manifest via an array of symptoms including claudication, ischemic rest pain, gangrene, and non-healing ulcers. Conventional risk factors for the development of PAD include smoking, age, hypertension, dyslipidemia, diabetes, and physical activity (4). Several non-conventional risk factors including inflammation, HIV, and exposure to toxic metals and air pollution have also emerged. Additionally, an underappreciated risk factor that increases the risk of developing PAD is chronic kidney disease (CKD) (5). Recently studies have estimated that approximately 25-35% of CKD patients develop PAD (1, 2, 6). A meta-analysis of 44,138 patients demonstrated that PAD patients with CKD have a ∼2-fold increase in major limb amputation and ∼2.55-fold increase in mortality risk (7).

These observations agree with an earlier examination of a large cohort of male veterans which reported that PAD patients with renal insufficiency were more likely to present with ischemic ulceration/gangrene and had greater mortality rates compared to PAD patients with normal renal function (8). Unfortunately, the success rates of endovascular and surgical interventions are significantly lower in PAD patients with CKD compared to those with normal kidney function (1, 8–10). Despite the widespread prevalence of renal insufficiency in PAD patients and worsened clinical outcomes, most clinical trials investigating novel therapeutics in PAD exclude patients with CKD. Thus, there is a significant gap in knowledge in understanding the coalescence of these diseases.

Although an association between CKD and PAD exists, the mechanisms linking the two pathologies are ill-defined. Independent of PAD, CKD has also been shown to negatively impact the vasculature, including a reduction in capillary density in the heart (11) and skeletal muscle (12, 13). Additionally, rodents with CKD have impaired angiogenic responses to ischemia (14–16), but mechanistic data to show how these impairments arise remains limited. Other factors that contribute to PAD pathogenesis and worsen limb outcomes include microvascular disease (17, 18), decreased vasodilatory capacity (19, 20) and increased inflammatory response in the endothelial cells (21), all of which may be exacerbated in patients with renal insufficiency. The retention of uremic solutes in the blood and tissues is a hallmark of CKD (22) and a recent study in PAD patients reported that plasma concentrations of tryptophan-derived uremic solutes were strongly associated with an increased risk of a major adverse limb event (16). Most notably, plasma concentrations of indoxyl sulfate, kynurenine, and kynurenic acid had hazard ratios of 2.2, 4.2, and 2.5 respectively. Reports in the literature have provided evidence that indoles and kynurenine metabolites (which accumulate in CKD) can impair nitric oxide-dependent vasodilation and angiogenesis (23–26), but the molecular mechanisms are incompletely understood. Several tryptophan-derived uremic metabolites (27–29) have been identified as ligands for the aryl hydrocarbon receptor (AHR), a ligand-activated transcription factor involved in xenobiotic metabolism. Elevated expression of the AHR is present in humans and rodents with CKD (30). Activation of the AHR using high affinity ligand 2,3,7,8-tetrachlorodibenzo-p-dioxin has been shown to promote atherosclerosis (31, 32). AHR has also been reported to regulate angiogenesis, although there are discrepancies in the literature with some studies reporting anti- (16, 33–35) and others reporting pro-angiogenic effects (36, 37) suggesting that these effects could be tissue and ligand specific. However, in a recent study where female mice with CKD were subjected to hind limb ischemia, treatment with an AHR inhibitor (CH223191) improved perfusion recovery and capillary densities in the ischemic limb (16). In this study, we sought to determine the impact of AHR activation, specifically in endothelial cells, on ischemia-induced angiogenesis and myopathy in mice with CKD. It was hypothesized that an endothelial cell-specific conditional deletion of the AHR would improve ischemic limb outcomes in mice with CKD.

## RESULTS

### Endothelium-Specific AHR Deletion Promotes Ischemic Perfusion Recovery and Alters Angiogenic Associated Gene Expression in Male Mice with CKD

To begin to explore whether chronic AHR activation in the endothelium contributes to worsen limb outcomes in PAD, we generated a conditional endothelial cell-specific AHR knockout mouse (AHR^ecKO^, **Figure 1A**). Following administration of tamoxifen, deletion of the AHR was confirmed by traditional PCR designed to detect recombination of exon 2 in the AHR gene (**Figure 1B**) as well as qPCR analysis of *Ahr* expression (**Figure 1C**), both performed in primary endothelial cells.

**Figure 1.**
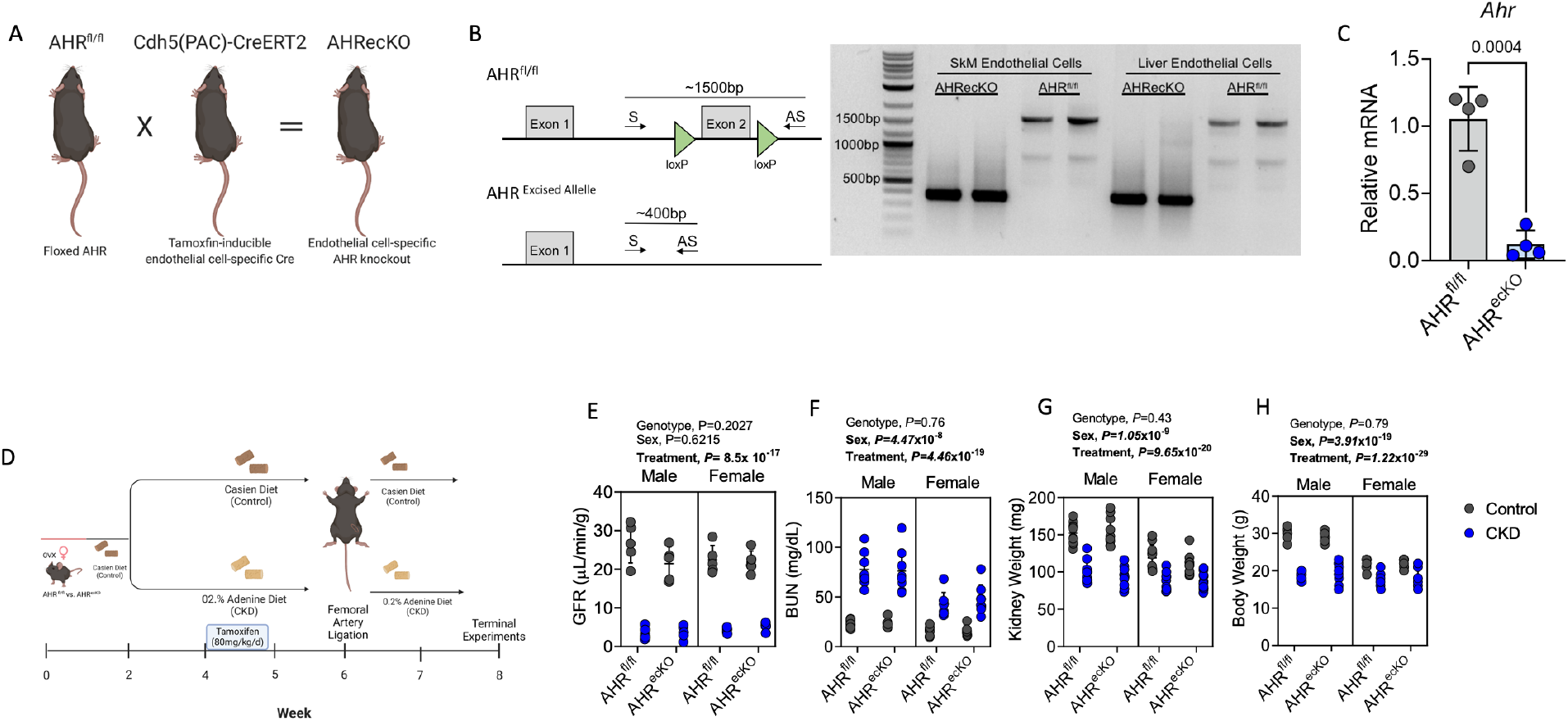
Validation of Endothelium-Specific Knockout of the AHR. (A) Generation of inducible, endothelial cell-specific AHR knockout mice. (B) Confirmation of DNA recombination in endothelial cells isolated from liver and skeletal muscle. (C) Confirmation of AHR deletion by qPCR analysis of mRNA expression. (D) Graphical depiction of the overall experiment design. (E) Glomerular filtration rate (GFR) measured by FITC-inulin clearance and normalized to body weight (n = 5/group/sex). (F) Blood urea nitrogen (BUN) measured in serum after eight weeks on adenine diet (n = 8/group/sex). (G) Kidney weights and (H) body weights measured at sacrifice. Panel C was analyzed using unpaired, two-tailed Student’s *t*-test. Panels E-H were analyzed using a three-way ANOVA. Error bars represent the standard deviation.

Additionally, endothelial-specific Cre-induced DNA recombination was confirmed using a fluorescent reporter mouse (**Supplemental Figure 1**; Ai14, Jackson laboratory, Stock #007914). Next, mice were randomized to either a casein (control) or adenine-supplemented (CKD) diet prior to the induction of limb ischemia via femoral artery ligation (FAL) surgery (**Figure 1D**).

Consistent with previous studies (38–40), mice fed adenine diet had lower glomerular filtration rate (GFR, **Figure 1E**) and elevated blood urea nitrogen (BUN, **Figure 1F**), which was accompanied by significantly lower kidney (**Figure 1G**) and body (**Figure 1H**) weights compared with mice fed the casein diet.

Because previous work has implicated that AHR activation can impair angiogenesis (16, 41, 42), we first examined whether deletion of the AHR in endothelial cells alone would alter perfusion recovery following FAL in mice with and without CKD. In the paw, there was a significant effect of diet in both male and female mice, indicating that CKD impaired perfusion recovery (**Figure 2A**). Interestingly, in the gastrocnemius muscle, the effect of diet was present only in male mice (**Figure 2B**), suggesting that CKD was more detrimental to males than females. A significant Time x Diet x Genotype interaction was detected (*P*=0.0146) for perfusion recovery in the paw of male mice. Post-hoc testing revealed significant higher levels of paw perfusion in male AHR^ecKO^ mice with CKD compared with male AHR^fl/fl^ mice with CKD at day 7 (*P*=0.0195) and day 14 (*P*=0.0002) after FAL. However, an interaction was not observed in female mice demonstrating that deletion of the AHR in endothelial cells had no impact on paw perfusion recovery in female mice. Similar to the paw, gastrocnemius muscle perfusion recovery was greater in male AHR^ecKO^ mice with CKD compared with male AHR^fl/fl^ mice with CKD at day 3 (*P*=0.0001) and day 7 (*P*=0.0227) after FAL, however, no difference was detected in female mice (**Figure 2B**).

**Figure 2.**
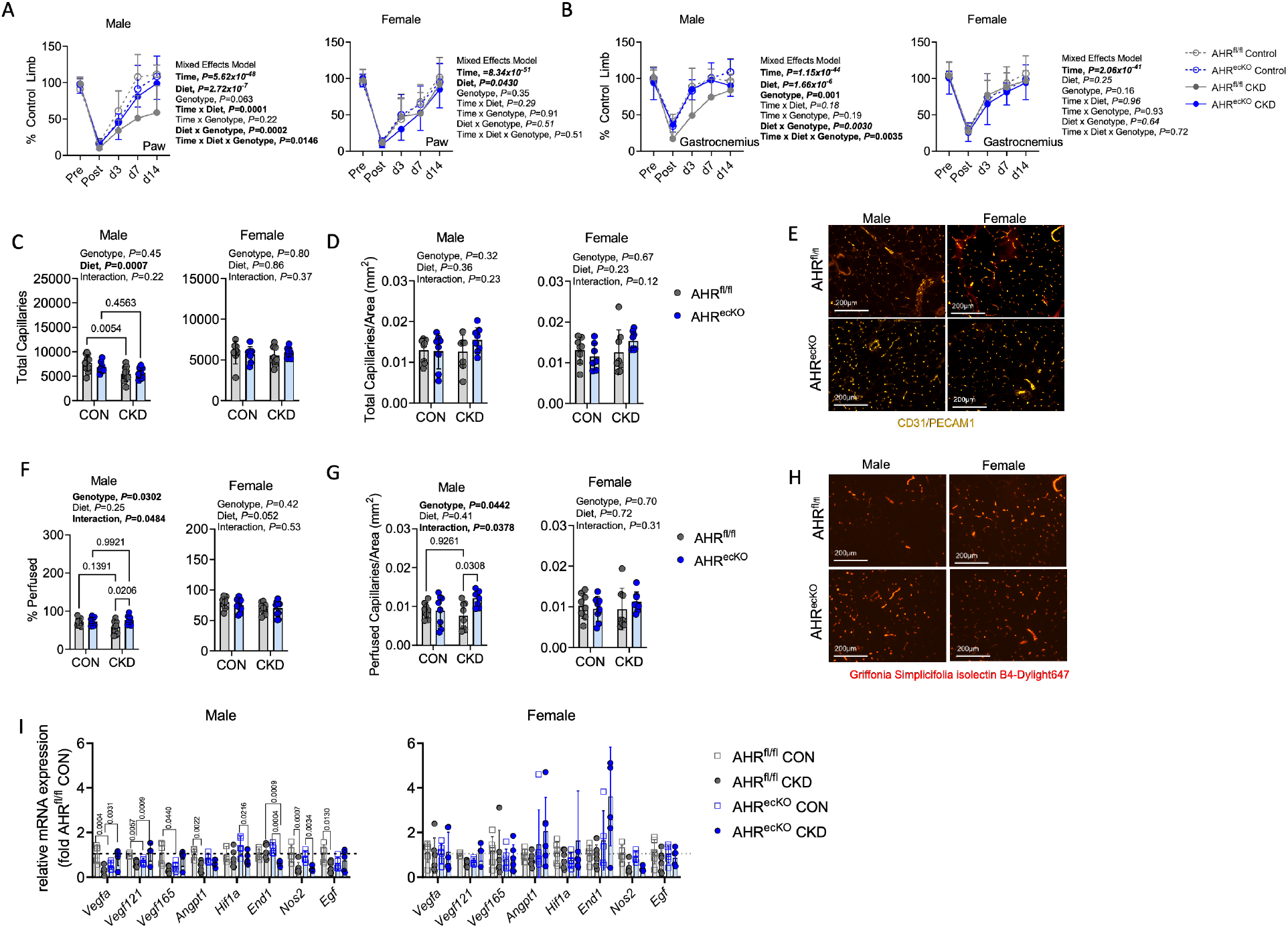
Endothelium-Specific AHR Deletion Promotes Ischemic Perfusion Recovery and Alters Angiogenic Associated Gene Expression in Male Mice with CKD. (A) Laser Doppler flowmetry quantification of perfusion recovery in the paw and (B) gastrocnemius muscle expressed as percentage of control limb (n=10/group). Perfusion recovery was analyzed using mixed model analysis. (C) Total number of capillaries and (D) total number of capillaries normalized to total muscle area. (E) Representative immunofluorescence images of total capillaries labeled with the anti-CD31 antibody. (F) Perfused capillaries quantified as a percentage of total capillaries. (G) Perfused capillaries quantified as the number per area of the muscle. (H) Representative immunofluorescence images of perfused capillaries labeled with isolectin. All capillary density measurements were performed n=8-10/group/sex. (I) Expression of angiogenic and vasoreactive genes in ischemic skeletal muscle (n=6/group/sex). Statistical analyses performed using two-way ANOVA with Sidak’s post hoc testing for multiple comparisons when significant interactions were detected. Error bars represent the standard deviation.

Because of the superficial penetration depth of the laser Doppler, we also performed a retro-orbital injection of Dylight649-labeled isolectin to live mice allowing for labeling and quantification of both perfused (isolectin^+^) and total (CD31^+^) capillaries across the tibialis anterior muscle. In male mice, CKD resulted in a significant decrease in the total capillary number in AHR^fl/fl^ mice only (**Figure 2C**), however, once normalized to the total muscle area this difference was abolished (**Figure 2C**). In female mice, total capillary number and total capillaries per area of muscle were unaffected by CKD or deletion of the AHR in endothelial cells (**Figure 2D**). In agreement with the laser Doppler findings, the percentage of perfused capillaries, as well as the number of perfused capillaries per muscle area was significantly higher in male AHR^ecKO^ mice with CKD compared to AHR^fl/fl^ mice with CKD **(Figure 2F)**. However, this relationship was not present in the female mice (**Figure 2G**). No significant differences in total or perfused capillaries were observed in the non-ischemic limb (**Supplemental Figure 2A,B**). Next, we employed qPCR to evaluate how the endothelial specific AHR deletion affects expression of angiogenic and vasoreactive genes in ischemic skeletal muscle (**Figure 2I**). In males, CKD resulted in decreased expression of pro-angiogenic genes *Vegfa* (*P*=0.0004), *Vegf121* (*P*=0.0057*), Vegf165* (*P*=0.0059), *Angpt1* (*P*=0.0022), and *Egf* (*P*=0.0130) in AHR^fl/fl^ mice. In contrast, only *Vegf121* was significantly reduced by CKD in AHR^ecKO^ male mice. In fact, AHR^ecKO^ mice with CKD had significantly higher expression of *Vegfa* (*P*=0.0031) and *Vegf121* (*P*=0.0009), along with a trending increase in *Vegf165* (*P*=0.0778), when compared to AHR^fl/fl^ male mice with CKD. In addition to angiogenic genes, the expression of endothelin 1 (*End1*), a vasoconstricting gene, was significantly reduced in male AHR^ecKO^ mice with CKD when compared to their AHR^fl/fl^ littermates (**Figure 2I**). Neither CKD nor the deletion of AHR were found to have significant impact on angiogenic or vasoreactive gene expression in female mice with or without CKD (**Figure 2I**). In the non-ischemic limb, the expression of angiogenic and vasoreactive genes displayed greater impact in males compared to females, but minimal effects of AHR deletion were observed (**Supplemental Figure 2C,D**).

### Endothelium-Specific AHR Deletion Does Not Improve Ischemic Muscle Function

Skeletal muscle function has emerged a critically important characteristic in PAD and has been shown to predict/associate with mortality risk and mobility loss (43–46). Additionally, independent of PAD, CKD is known to manifest with a progressive skeletal myopathy characterized by muscle atrophy, weakness, and exercise intolerance (47, 48). Thus, we tested whether deletion of the AHR in endothelial cells influenced ischemic muscle function. First, we used voluntary unilateral hindlimb grip strength testing to evaluate paw function and strength. There was a significant main effect of CKD (diet) in both male (*P*=2.72E-07) and female mice (*P*=0.0430) demonstrating that CKD decreased voluntary grip strength in the ischemic limb (**Figure 3A**). A significant Time x Genotype x Diet interaction was detected only in male mice. However, post-hoc testing did not reveal any significant differences in grip strength between AHR^fl/fl^ and AHR^ecKO^ male mice (*P*=0.16 at day 14 post-FAL). To provide a more rigorous assessment of the ischemic limb muscle function, we utilized nerve mediated contraction of the tibialis anterior *in-situ*, where supramaximal stimulation could be applied to ensure complete recruitment of the motor neurons. Consistent with previous findings in CKD mice subjected to FAL (39), male mice with CKD had lower twitch (**Figure 3B**) and tetanic (**Figure 3C**) forces, regardless of whether peak force was normalized to the muscle weight or not (**Figure 3D-G**). Despite the improved perfusion recovery and increased perfused capillary density in male AHR^ecKO^ mice with CKD, muscle contractile function was not different from AHR^fl/fl^ mice (**Figure 3D-G**). In female mice, CKD did not significantly impair muscle contractile function following FAL and consequently the deletion of AHR in endothelial cells had no significant impact on muscle strength (**Figure 3H-M**). In the non-ischemic limb of mice with CKD, deletion of the AHR in endothelial cells had no effect on muscle contractile function in either sex (**Supplemental Figure 3**).

**Figure 3.**
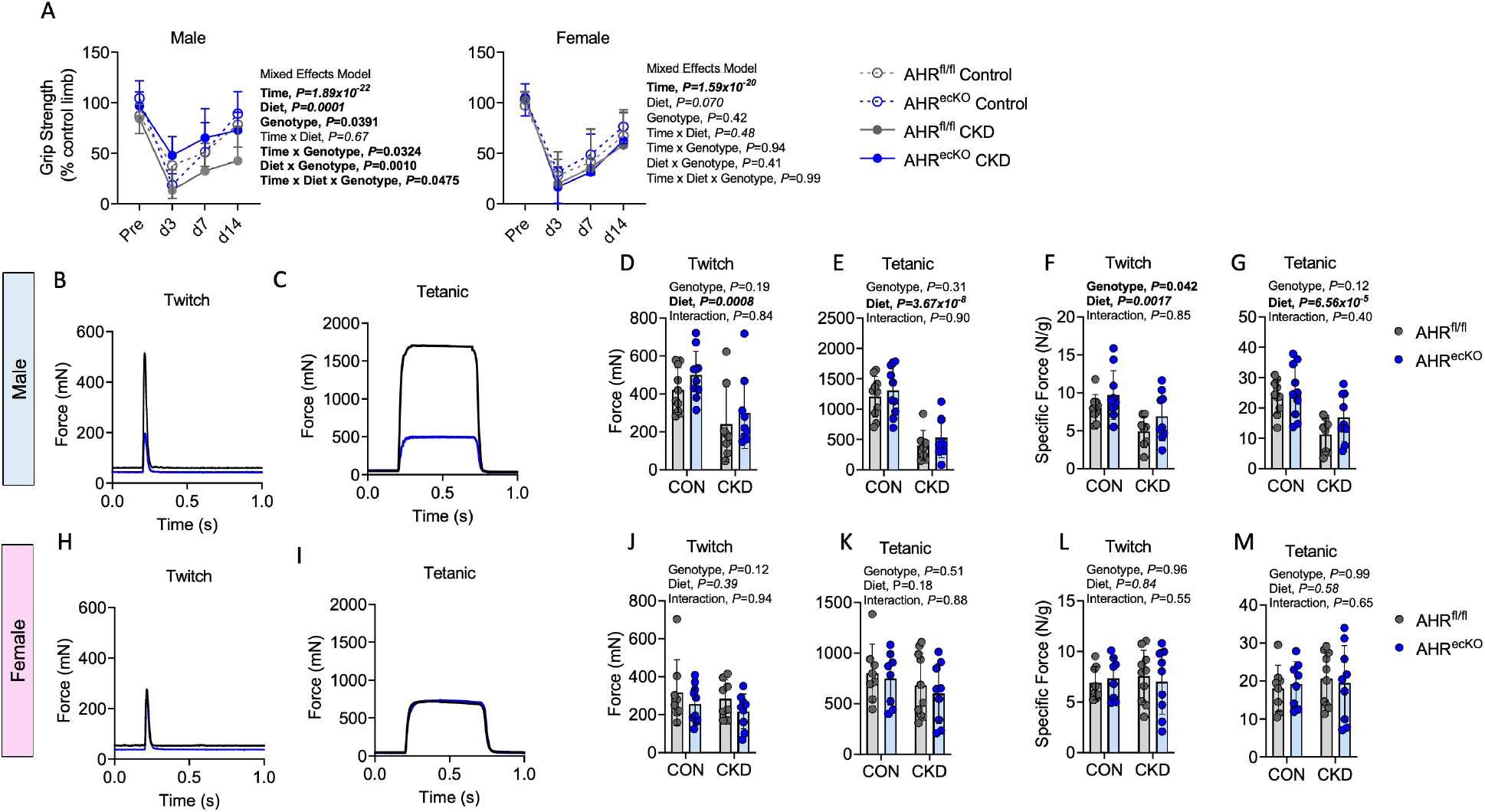
Endothelium-Specific AHR Deletion Does Not Improve Ischemic Muscle Function. (A) Hind limb grip strength recovery measured in grams force and quantified as percentage of control limb. Grip strength recovery was analyzed using mixed model analysis (n=10/group/sex). (B) Representative twitch and (C) tetanic contractions in male mice with and without CKD. (D) Maximal absolute twitch force, (E) maximal absolute tetanic force, (F) maximal specific (normalized to weight) twitch force, (G) maximal specific tetanic force in male mice. (H) Representative twitch and (I) tetanic contractions in female mice with and without CKD. (J) Maximal absolute twitch force, (K) maximal absolute tetanic force, (L) maximal specific (normalized to weight) twitch force, (M) maximal specific tetanic force in female mice. Panels D-M were analyzed using two-way ANOVA with Sidak’s post hoc testing for multiple comparisons. Error bars represent the standard deviation. Panels D-G and J-M contain n=8- 10/group.

### Endothelium-Specific AHR Deletion Has No Impact on Muscle Mass or Histopathology but Increases Myofiber Area in Male CKD Mice

Next, we examined the impact of CKD and endothelium specific AHR deletion on muscle size, myofiber area, and muscle histopathology in the ischemic limb. There was a significant effect of CKD on the weight of the tibialis anterior and gastrocnemius muscles in both male and female mice, however there was no significant main effect for genotype or interaction detected (**Figure 4A,B**). Quantification of the total myofiber number within the tibialis anterior muscle revealed a significant impact of CKD in male mice only (*P*=0.0268), but no significant effects of genotype (**Figure 4C**). Representative images of ischemic muscle histopathology are shown in **Figure 4D**. There was no significant difference in percentage of myofibers with centralized nuclei among any of the groups (**Figure 4E**). Analysis of the mean myofiber cross-sectional area (CSA) revealed significantly smaller myofiber areas in male CKD mice (diet effect, *P*=1.9E-05), but not female mice (diet effect, *P*=0.27). Interestingly, male AHR^ecKO^ mice with CKD were found to have significantly higher mean myofiber CSA compared to AHR^fl/fl^ mice with CKD (**Figure 4F**). In the non-ischemic limb, CKD mice had lower muscle weights and myofiber CSA, but there were no significant effects of AHR deletion in mice with or without CKD regardless of biological sex (**Supplemental Figure 4**).

**Figure 4.**
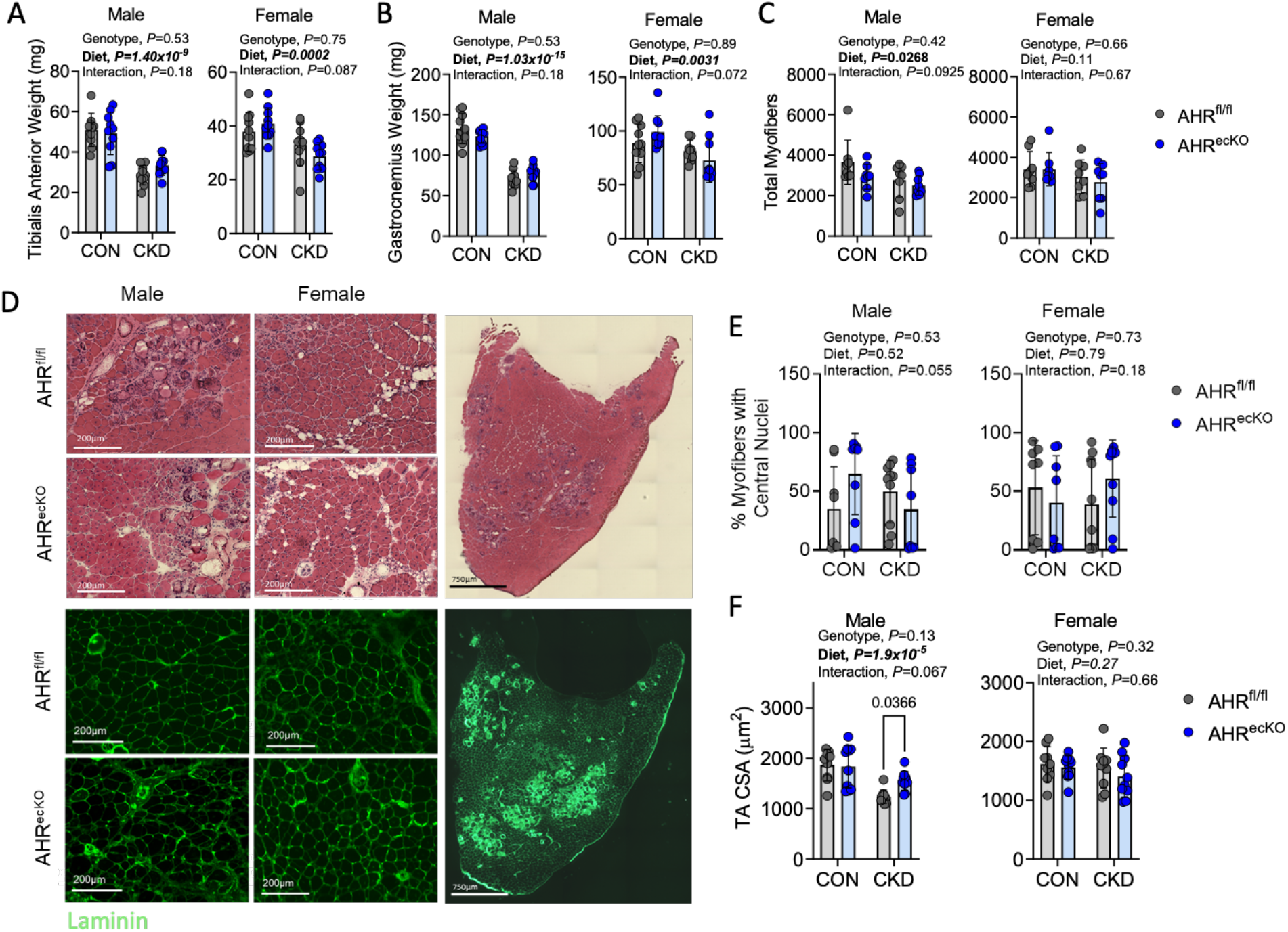
Endothelium-Specific AHR Deletion Has No Impact on Muscle Mass or Histopathology but Increases Myofiber Area in Male CKD Mice. (A) weight of tibialis anterior and (B) gastrocnemius muscles of the ischemic limb (n=10/group/sex). (C) Total number of myofibers within the ischemic tibialis anterior muscle (n=8/group/sex). (D) Representative 20x and tiled images of hematoxylin & eosin staining and laminin immunolabeling of ischemic tibialis anterior muscle. (E) Percentage of total myofibers with centralized nuclei (n=8-10/group/sex). (G) Quantification of mean myofiber cross-sectional area (CSA) of the ischemic tibialis anterior muscle (n=10/group/sex). Statistical analyses performed using two-way ANOVA with Sidak’s post hoc testing for multiple comparisons when significant interactions were detected. Error bars represent the standard deviation.

### Endothelium-Specific AHR Deletion Does Not Impact Ischemic Muscle Mitochondrial Function in CKD

Alterations in mitochondrial respiration and energy production have been identified as characteristics of both skeletal muscle in both PAD and CKD independent of one another (38, 49–56). Thus, we sought to examine if deletion of the AHR in endothelial cells and the corresponding improvement in limb perfusion recovery in male AHR^ecKO^ mice altered skeletal muscle mitochondrial oxidative phosphorylation (OXPHOS). To accomplish this, we employed a creatine kinase clamp system which facilitates the assessment of mitochondrial oxygen consumption (*J*O_2_) at various energy demands, akin to a stress test. Quantification of the slope of the relationship between *J*O_2_ and energy demand (ΔG_ATP_), termed OXPHOS conductance, represents the mitochondrion’s ability to respond to changes in energy demand by altering energy transduction. For these experiments, mitochondria were isolated from the ischemic gastrocnemius muscle and energized with saturating levels of pyruvate, malate, and the medium chain fatty acid octanoylcarnitine (**Figure 5A**). Graphs of the *J*O_2_ and ΔG_ATP_ relationship in for male and female mice are shown in **Figure 5B and 5C**. Quantification of the OXPHOS conductance revealed a significant genotype effect in male mice (*P*=0.0021) only (**Figure 5C**). Posthoc analysis revealed that CON male AHR^ecKO^ mice had higher OXPHOS conductance than CON AHR^fl/fl^ mice (*P*=0.0316), but there was no difference between genotypes in the CKD group (*P*=0.45, **Figure 5D**). The respiration rate at the highest level of energy demand was also significantly higher in CON AHR^fl/fl^ mice (*P*=0.0265), but there was no difference between genotypes in the CKD group (*P*=0.25, **Figure 5E**). Mitochondrial hydrogen peroxide (H_2_O_2_) production and an estimation of electron leak were not affected by CKD or genotype in male mice (**Figure 5F,G**). In female mice, there was a significant diet effect indicating CKD mice had lower OXPHOS conductance (**Figure 5H**) and respiration rates at the highest level of energy demand, but no genotype effect was observed (**Figure 5I**). Mitochondrial hydrogen peroxide (H_2_O_2_) production and an estimation of electron leak were not affected by CKD or genotype in female mice (**Figure 5J,K**). In the non- ischemic limb, deletion of the AHR in endothelial cells did not impact muscle mitochondrial function in mice with or without CKD (**Supplemental Figure 5**). In the ischemic muscle, the transcript levels of *Cox7a1*, *Atp5k*, *Atp5d*, *Tfam*, and *Sod2* were reduced by CKD in AHR^fl/fl^ (**Figure 5L**). Interestingly, male AHR^ecKO^ mice without CKD tended to have lower expression of these mitochondrial genes compared to AHR^fl/fl^ mice without CKD, but the impact of CKD in AHR^ecKO^ was smaller and only *Tfam* expression was significantly decreased by CKD in male AHR^ecKO^ mice. Female mice did not have significant changes in mitochondrial gene expression in the ischemic limb due to either CKD or genotype (**Figure 5M**). However, in the non-ischemic limb, CKD was found to alter the expression of several of these mitochondrial genes in both sexes, but a genotype effect was only observed in a few mitochondrial genes with AHR^ecKO^ mice on the control diet having lower expression compared to AHR^fl/fl^ without CKD (**Supplemental Figure 5**).

**Figure 5.**
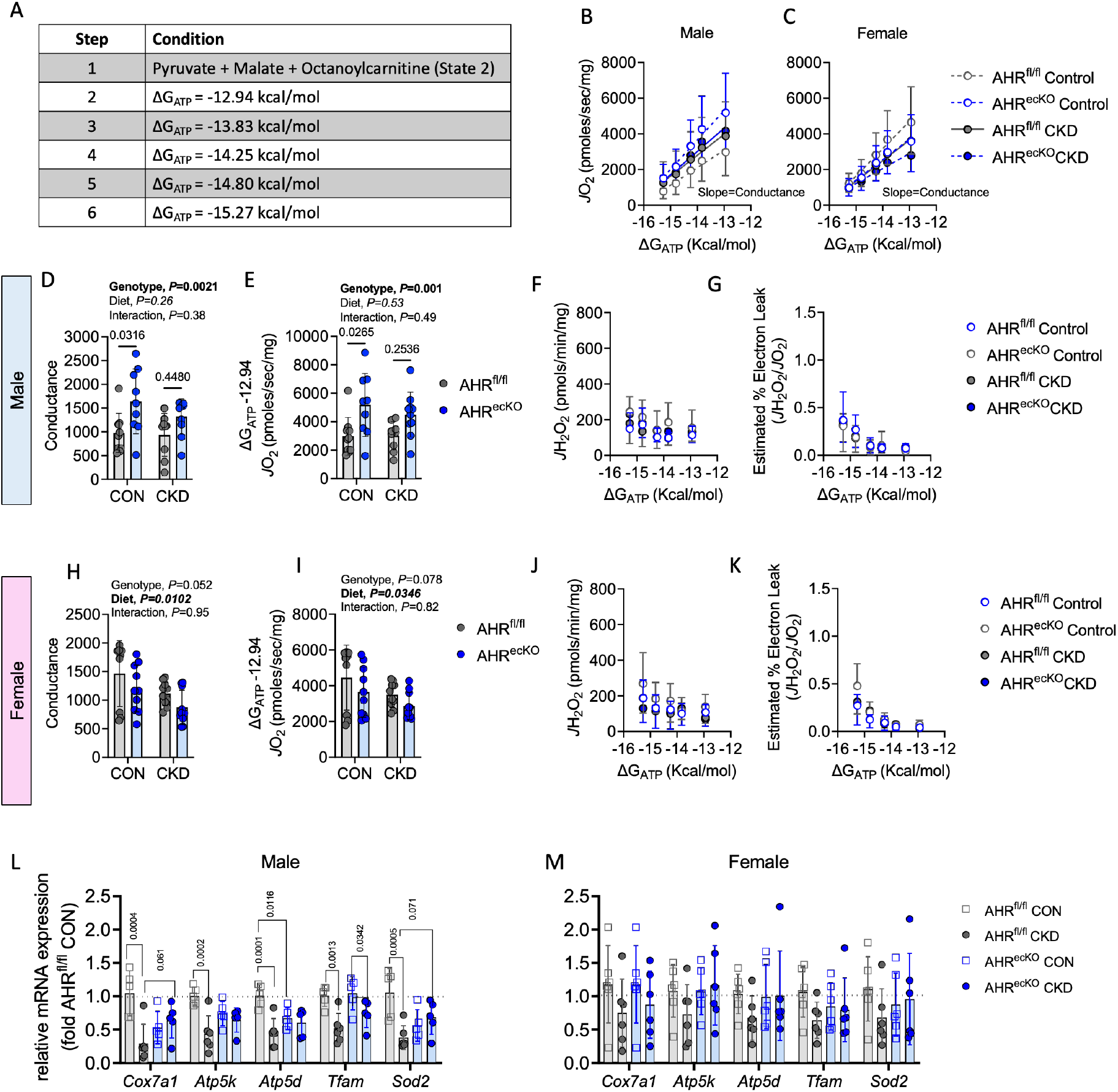
Endothelium-Specific AHR Deletion Does Not Impact Ischemic Muscle Mitochondrial Function in CKD: (A) Protocol for assessing mitochondrial function in the ischemic gastrocnemius muscle was assessed using a creatine kinase clamp system to measure oxygen consumption (*J*O_2_) at physiologically relevant energy demands (ΔG_ATP_). (B) Plot of *J*O_2_ and ΔG_ATP_ in mitochondria isolated from male and (C) female mice. (D) Quantification of the conductance (slope of *J*O_2_ and ΔG_ATP_ relationship) and (E) *J*O_2_ at the highest energy demand in male mice. (F) Mitochondrial H_2_O_2_ production and (G) estimated percent electron leak (*J*H_2_O_2_/ *J*O_2_) at each ΔG_ATP_ in males. (H) Quantification of the conductance (slope of *J*O_2_ and ΔG_ATP_ relationship) and (I) *J*O_2_ at the highest energy demand female mice. (J) Mitochondrial H_2_O_2_ production and (K) estimated percent electron leak (*J*H_2_O_2_/ *J*O_2_) in females. (L) expression of mitochondrial related genes in ischemic extensor digitorum longus muscle in male and (M) female mice (n=6/group/sex). Statistical analyses performed using two-way ANOVA with Sidak’s post hoc testing for multiple comparisons when significant interactions were detected. Error bars represent the standard deviation. Panels B-K contain n=10/group/sex.

### Endothelial Cell AHR Displays Sex-Dependent Differences in Activating Potential in Normoxic and Hypoxic Conditions

An intriguing observation in this study was a sex-dependent effect of AHR activation on ischemic angiogenesis. While the underlying mechanisms behind this sexual dimorphism in endothelial cells are unknown, sex-dependent effects of AHR activation have been shown in other tissues (57–60). To determine if these differences were intrinsic to biological sex and not dependent on sex hormones, we isolated primary endothelial cells from male and female mice, cultured them *ex vivo* where sex hormones were not present, and exposed cells to the potent AHR ligand indoxyl sulfate (IS) under normoxic and hypoxic conditions (**Figure 6A**). The purity of primary endothelial isolates was confirmed by immunolabeling with an anti-CD31 antibody (**Figure 6B**). Following treatment with indoxyl sulfate or DMSO (vehicle control) for six hours, primary endothelial cells were harvested for qPCR analysis of mRNA levels of the *Ahr*, *Cyp1a1*, and the *Ahrr* (AHR repressor). *Ahr* expression displayed a significant main effect for hypoxia and a hypoxia x indoxyl sulfate interaction (**Figure 6C**). Posthoc testing revealed a ∼12-fold increase in *Ahr* expression in hypoxic cells treated with indoxyl sulfate in both male and female primary endothelial cells (**Figure 6C**). To examine the activating potential of the AHR, we measured expression of *Cyp1a1*, a canonical xenobiotic response gene whose expression is controlled by *Ahr*. As expected, there was a main effect of indoxyl sulfate (*P*=0.0001) fully demonstrating that indoxyl sulfate treatment upregulates *Ahr* signaling. Interestingly, a significant main effect of sex, as well as an indoxyl sulfate x sex interaction was detected. Posthoc analysis revealed higher *Cyp1a1* expression levels in male primary endothelial cells treated with indoxyl sulfate when compared to female primary endothelial cells under both normoxic and hypoxic conditions (**Figure 6D**).

**Figure 6.**
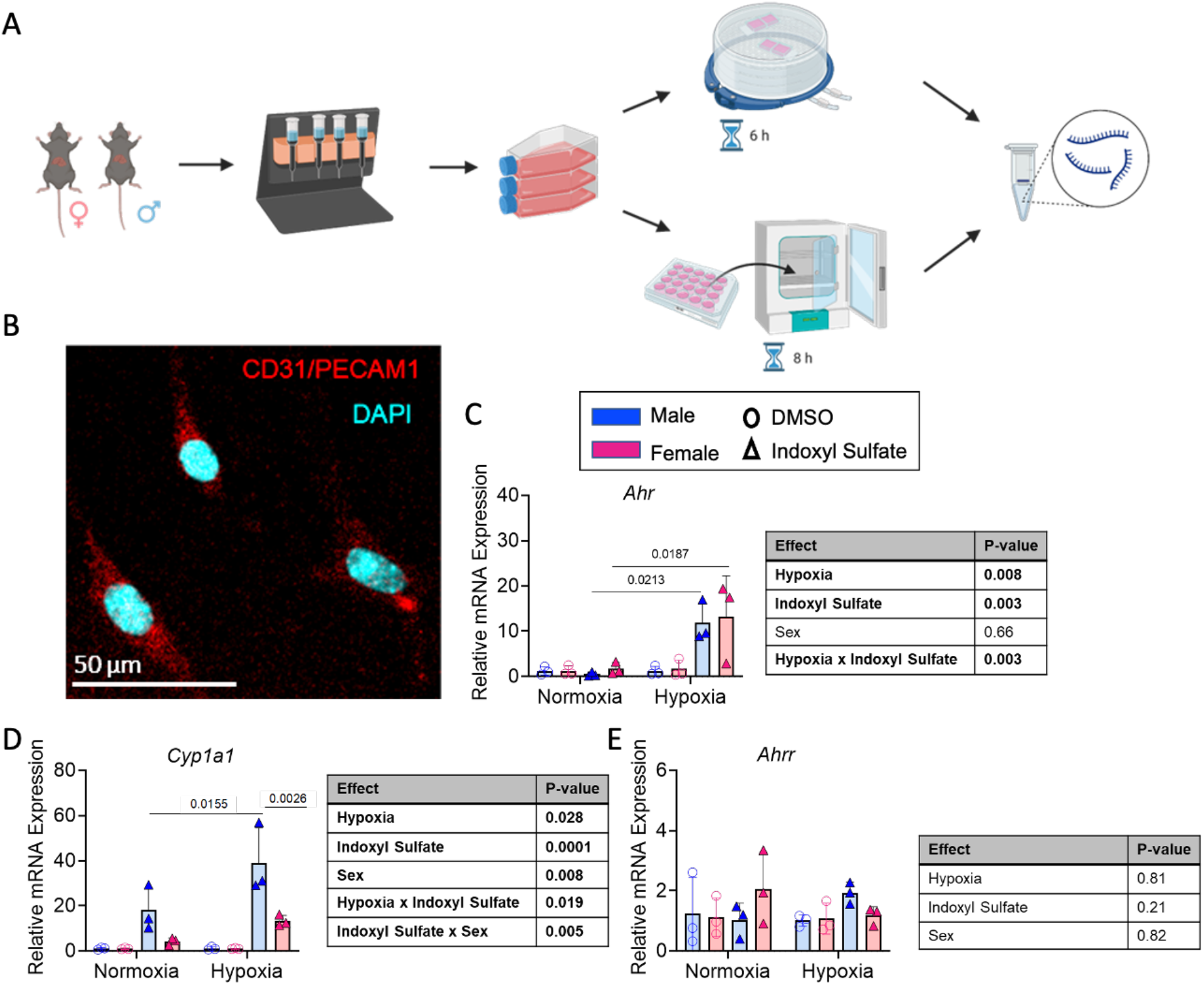
Endothelial Cell AHR Displays Sex-Dependent Differences in Activating Potential in Normoxic and Hypoxic Conditions: (A) Graphical depiction of experimental design. (B) Confirmation of endothelial isolates via immunolabeling with an anti-CD31 antibody. (C) Relative mRNA expression of *Ahr* (D) *Cyp1a1* and (E) *Ahrr* measured by qPCR analysis in primary cells treated with indoxyl sulfate or DMSO control in normoxic and hypoxic conditions (n=3 biological replicates/group/sex). Statistical analyses performed using two-way ANOVA with Sidak’s post hoc testing for multiple comparisons when significant interactions were detected. Error bars represent the standard deviation.

The determine if the lower AHR activating potential in female endothelial cells was due to elevated expression of the AHR repressor system, we measured *Ahrr* expression levels.

Importantly, *Ahrr* expression was found to be unaffected by either sex or hypoxia (**Figure 6E**). Taken together, these findings demonstrate that male endothelial cells display greater AHR activating potential when compared to female endothelial cells, and this difference is not explained by repression of the AHR or differences in sex hormones.

### RNA Sequencing in Primary Endothelial Cells Uncovers Sex-Dependent Alterations in Angiogenic Signaling with AHR Activation in Hypoxia

To assess the broader impact of sex-specific effects on *Ahr* activation on the endothelial cell transcriptome, we performed RNA sequencing on male and female primary endothelial cells treated with indoxyl sulfate under conditions of hypoxia and nutrient deprivation. There were 1076 upregulated (Log_2_FC < -1.5, adjusted *P*<0.05) and 690 downregulated (Log_2_FC > 1.5, adjusted *P*<0.05) in male compared to female endothelial cells (**Figure 7A**). A complete list of differentially expressed genes can be found in Supplemental Dataset 1. Gene set enrichment analysis of genes upregulated in male endothelial cells revealed terms related to *“morphogenesis of a branching structure”*, *“regulation of angiogenesis”*, *“wound healing”*, *“epithelial tube morphogenesis”,* and *“ameboidal-type cell migration”* (**Figure 7B**). A heatmap of the top 100 differentially expressed genes is shown in **Figure 7C**. Despite being exposed to hypoxia and nutrient deprivation within the same culture plate, *Hif1a* expression ∼4.6-fold higher in male endothelial cells compared to female endothelial cells (adjusted *P*=0.0057). Correspondingly, several genes involved in angiogenesis were also expressed differently in the sexes. For example, males had higher expression of *Vegfa, Vegfb,* and *Hgf,* whereas females had higher expression of *Fgf2* (**Figure 7D**). These data also exposed sex-dependent differences in the Notch signaling pathway, a master regulator of sprouting angiogenesis (61), under conditions of *Ahr* activation. Male endothelial cells had greater expression of *Notch1/4* and notch-related genes *Dll4* and *Cxcr4*, as well as decreased expression of *Usp10*, an antiangiogenic factor that inhibits Notch (62). Male endothelial cells also had higher expression of matrix metalloproteinases (*Mmp2/9/19*) which are involved in remodeling of the extracellular matrix during vessel sprouting (63). Several genes that regulate redox balance and have been linked to angiogenesis (64) also displayed sex differences with males expressing less *Trx1*, *Trxnd1*, but more *Txnip* and *Sod3* when compared to female endothelial cells (**Figure 7D**). Male endothelial cells also had greater expression of nitric oxide synthase 3 (*Nos3*) compared to female endothelial cells (adjusted *P*=2.8E-04) treated with indoxyl sulfate, suggesting that vasoreactivity may also be different between males and females with AHR activation.

**Figure 7.**
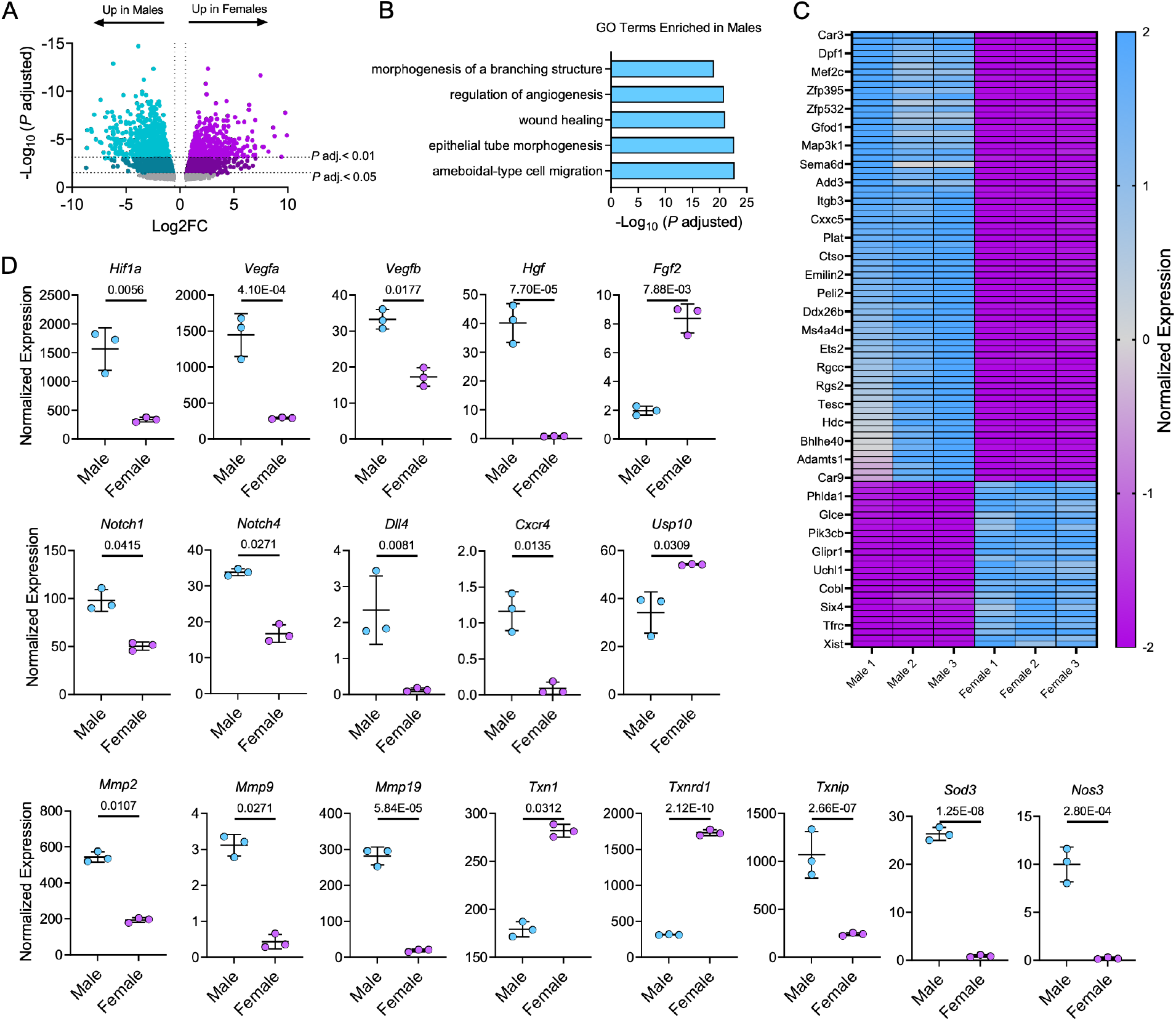
RNA Sequencing in Primary Endothelial Cells Uncovers Sex-Dependent Alterations in Angiogenic Signaling with AHR Activation in Hypoxia: (A-D) RNA extracted from primary endothelial cells isolated from male and female mice treated with indoxyl sulfate under hypoxic conditions. (A) Volcano plot of all detectable genes, differential expression significant if (Log_2_FC < -1.5, adjusted *P*<0.05) or (Log_2_FC > 1.5, adjusted *P*<0.05). (B) Gene set enrichment analysis of genes upregulated in male endothelial cells. (C) Heat map of top 100 differentially expressed genes. (D) Normalized expression of select differentially expressed genes from top GO terms. Adjusted *P*-values from a Wilcoxon test are shown in Panel D.

## DISCUSSION

The presence of CKD is known to exacerbate the pathobiology of PAD and increase the risk of major adverse limb events and mortality. Despite the robust clinical evidence, the cellular mechanisms that contribute to worsened outcomes are ill defined. In this study we utilized adenine diet and femoral artery ligation to explore the role of AHR activation, caused by uremic solute accumulation, in regulating ischemic angiogenesis in CKD. In male mice with CKD and PAD, endothelial cell-specific deletion of the AHR improved limb perfusion recovery, enhanced angiogenesis evidenced by more perfused capillaries, and attenuated myofiber atrophy. Gene expression analysis revealed greater mRNA levels of angiogenic and mitochondrial genes within the ischemic limb of AHR^ecKO^ mice with CKD compared to AHR^fl/fl^ mice with CKD. However, these effects did not translate to improvements in muscle strength or mitochondrial function in male AHR^ecKO^ mice. Conversely, female AHR^ecKO^ mice with CKD had no significant improvement in limb perfusion recovery, angiogenesis, or muscle health. Taken together, these findings demonstrate that endothelial cell AHR activation diminishes ischemic angiogenesis in male but not female mice with CKD.

The results herein are in agreement with previous studies demonstrating that AHR activation impairs angiogenic processes in cultured endothelial cells (33, 65–67) and mice (14, 16). A recent study in mice with CKD provided evidence implicating activation of the AHR in the degradation of beta catenin leading to disruption of the Wnt signaling pathway which impaired ischemic angiogenesis (16). The authors also found that global AHR inhibition improved perfusion recovery and normalized ischemic angiogenesis in female mice with CKD. In contrast, the results herein demonstrated that genetic deletion of the AHR specifically in endothelial cells improved ischemic angiogenesis in male, but not female, mice with CKD. The discrepancy between our study and Arinze *et. al*. may be explained by differences in the experimental approach. First, while both studies used adenine-supplemented diet to induce CKD, the duration of CKD was significantly longer in the current study (6 vs. 3 weeks). Second, our study employed a genetic ablation of the AHR specifically in endothelial cells, whereas Arinze *et. al*. treated mice with CH223191, a pharmacologic AHR inhibitor. Recent evidence has established that AHR activation has pathological effects in skeletal muscle (68) which can disrupt paracrine angiogenic signaling (CITE Circ Res). Thus, it is likely that some of the observed benefits with CH223191 treatment were derived from the drug’s effects on limb muscle.

Unexpectedly, we found sex-specific differences when comparing the effect of AHR deletion in endothelial cells during CKD on ischemic limb pathology. Whereas male AHR^ecKO^ mice with CKD had improved limb perfusion recovery and angiogenesis, their female counterparts were not affected by AHR deletion. The notion that AHR signaling may be sex-dependent is supported by previous work in non-vascular tissue. For instance, treatment with the high-affinity AHR ligand, 2,3,7,8-tetrachlorodibenzo-p-dioxin, resulted in sex-differences in the liver transcriptome of mice (58). Additionally, in rodents with CKD, AHR abundance in the kidney has sexually dimorphic effects (57). Considering that all female mice underwent ovariectomies prior to enrollment in our study, we theorized that these stark differences may not be explained by sex hormones alone. To test this hypothesis, we isolated and cultured primary endothelial cells from male and female mice *ex vivo* where sex hormones were absent. Cells were treated with known AHR ligand and uremic solute, indoxyl sulfate, in normoxic and hypoxic conditions. These experiments confirmed that male endothelial cells had greater AHR activating potential in both normoxic and hypoxic condition compared to female endothelial cells. However, these differences were not related to the abundance of *Ahr* or its repressor (*Ahrr*) which were not different between male and females under these conditions. These data unequivocally establish that murine endothelial cells have sex-dependent differences in AHR activating potential which are not explained by sex hormones. To further explore this, we performed RNA sequencing on male and female primary hypoxic endothelial cells treated with IS. This experiment revealed sex-differences in several genes related to angiogenesis that agree with the observed sex-differences in ischemic angiogenesis in our mouse studies. In totality, these findings implicate sex chromosomes as potential mediator of the sex-dependent differences in AHR biology, however, future mechanistic studies are needed to confirm this assertion.

The current study along with previous work from our group (69) demonstrates that vascular targeted interventions can improve blood perfusion without impacting muscle function. Limb ischemia resulting from PAD induces angiogenesis, a process by which pre-existing endothelial cells proliferate and migrate to form new blood vessels (70, 71). Yet, patients with PAD have diminished microvascular perfusion (18, 21, 72–74) suggesting that the angiogenic response is insufficient to meet the demands of the tissue. Numerous studies have utilized gene, protein, and cell-based therapies aimed to promote angiogenesis in PAD patients, however these have been unsuccessful in clinical trials (75–81). While the lack of clinical success with angiogenic therapies is complex, it may indicate that targeting angiogenesis alone to improve PAD symptomology is suboptimal. It is important to recognize that it is challenging to adequately model the interaction between the PAD patients’ comorbidities, genetics, and environment in a preclinical setting. The poor modeling of PAD is likely to be a main contributor to the failure of translating preclinical PAD therapies to humans. It is indisputable that reduced blood flow contributes to walking performance in PAD patients, therefore, improving oxygen delivery to the limb should remain a primary clinical focus. However, there are several non-vascular cell types within the lower extremity (skeletal muscle, adipose tissue, connective tissue, nerves, etc.) that contribute to walking performance and how these cell type interact with vascular cells in the ischemic microenvironment is incompletely understood. Skeletal muscle can regulate angiogenesis via paracrine signaling (82) and coincidentally muscle quality/function in PAD patients is a strong predictor of morbidity and mortality (43–46). To make matters worse, common comorbidities in PAD patients like CKD have deleterious effects on skeletal muscle independent of PAD (47, 48, 83). In fact, several preclinical studies have demonstrated that therapeutic interventions targeting ischemic skeletal muscle can improve limb outcomes in mice subjected to femoral artery ligation (69, 84–87). The stagnant therapeutic development in PAD and lack of success of angiogenic therapies indicates that further research is necessary to understand the complex pathobiology of PAD and the intercellular communication contributing to limb functionality.

We acknowledge that there are limitations present in the current study. First, we enrolled young mice despite PAD prevalence being strongly associated with advancing age.

Unfortunately, due to the extensive breeding necessary to generate conditional, endothelial specific AHR deletion, experiments in aged mice were not possible. Second, limb ischemia in PAD patients occurs gradually over time whereas the FAL surgery used herein causes abrupt and severe ischemia. Additionally, mice were only studied at a single time-point post-FAL surgery such that temporal differences in angiogenic processes could not be evaluated. Third, PAD patients with CKD often present with additional comorbidities (hypertension, obesity, hyperlipidemia, diabetes) which contribute to disease pathology, but were not present in the mouse models employed in our study. Finally, while our study examined muscle histopathology and strength, our contraction protocol only measured muscle force levels during brief contractions. Thus, changes in muscle function may not translate the sustained contractions, such as the commonly used six-minute walk test in PAD patients. Future work is needed to develop better preclinical assessments of limb functionality as walking performance following a single-limb surgery (like FAL) is difficult to assess in quadrupedal species.

In summary, deletion of the AHR in endothelial cells improves limb perfusion recovery and ischemic angiogenesis in male, but not female, mice with PAD and CKD. However, the improved limb blood flow in males did not result in better muscle strength or mitochondrial function in the PAD limb. Primary endothelial cell culture experiments demonstrate sex-differences in AHR activation and angiogenic signaling that are not dependent on sex hormones.

## MATERIALS AND METHODS

### Animals

An endothelial cell-specific AHR knockout mouse (AHR^ecKO^) was generated by breeding floxed AHR mice (AHR^tm3.1Bra^/J, Jackson Laboratories, Stock No. 006203) with Cdh5(PAC)-Cre^ERT2^ mice (Taconic, Stock No. 13073). At 12 weeks of age mice, the conditional endothelial-specific AHR deletion was induced by delivering intraperitoneal injections of tamoxifen (80mg/kg) for five consecutive days. Littermates without the Cre transgene (AHR^fl/fl^) were injected with tamoxifen and used as controls. All experiments involved male and female mice. Female mice were ovariectomized two weeks prior to enrollment to mimic the sex hormone-deficient condition seen in post-menopausal PAD patients. Buprenorphine (0.05 mg/kg) was given post-operatively for analgesia. All mice were housed in temperature (22°C) and light-controlled (12:12-h light-dark) rooms and maintained on standard chow (Envigo Teklad Global 18% Protein Rodent Diet 2918 irradiated pellet) with free access to food and water prior to enrollment. All animal experiments adhered to the *Guide for the Care and Use of Laboratory Animals* from the Institute for Laboratory Animal Research, National Research Council, Washington, D.C., National Academy Press. To validate endothelium specific knockout of the *Ahr*, genomic DNA was extracted from CD31^+^ cells isolated from skeletal muscle and liver from AHR^fl/fl^ and AHR^ecKO^ mice using Qiagen DNeasy kit (Qiagen, Cat. No. 69504) according to manufacturer’s instructions. DNA was amplified using Terra PCR Direct Red Dye Premix (Takara Cat. No. 639286) and ran on a 1% agarose gel to confirm DNA recombination. Primer sequences are listed in Supplemental Table 1.

### Endothelial Cell Isolation

Primary CD31-positive cells were isolated from mouse liver and skeletal muscle as previously described (88). Briefly, transcardial perfusions with ice-cold phosphate-buffered saline (PBS) were performed via the left ventricle at a rate of 2mL/min for five minutes immediately followed by perfusion with digestion buffer (0.1% w/v collagenase type I, 0.1% w/v collagenase type II, 0.25U/mL Dispase, 7.5µg/mL DNAse in DMEM). Next, all hind limb skeletal muscles and the liver were dissected and minced separately on ice. The minced liver and skeletal muscle were then transferred separately into 15mL of digestion buffer and incubated at 37°C for 30 minutes. The digestion was stopped by adding 8mL of wash buffer (0.5% w/v bovine serum albumin and 2mM EDTA in PBS). Cell suspensions were then filtered through a 100µm cell strainer and centrifuged at 300xG for seven minutes. The supernatant was discarded, and the remaining cell pellet was resuspended in 5mL of wash buffer and centrifuged at 300xG for five minutes. The supernatant was discarded, and this step was repeated until the supernatant was clear. The final pellet was resuspended in 90µL of wash buffer and 10µL of CD31 Microbeads (Miltenyi Biotec, Cat. No. 130-097-418) were added to the suspension and incubated on ice for 20 minutes. Following microbead incubation, the suspension was washed, and the pellet was resuspended in 0.5mL of wash buffer. Suspension was passed through LS columns (Miltenyi Biotec, Cat. No. 130-042-401) on a magnetic separator (Miltenyi Biotec, Cat. No. 130-091-051). The columns were removed from the magnet and fraction containing CD31-positive cells were eluted using 5mL of wash buffer. CD31-positive fraction was centrifuged at 300xG for ten minutes, the supernatant discarded, and the remaining cell pellet was used for subsequent experiments.

### Induction of chronic kidney disease

All mice were fed a casein control diet for a seven-day acclimation period before being randomly assigned to remain on casein diet or to be fed a 0.2% adenine supplemented diet to induce CKD (38, 89, 90). Mice remained on their respective diet for four weeks prior to femoral artery ligation (FAL) and during the two-week post-FAL recovery period. All diets were provided ad libitum for the duration of the study.

### Assessment of kidney function

Kidney function was evaluated via glomerular filtration rate (GFR) and blood urea nitrogen (BUN) levels as previously described (38–40, 91–93). In brief, GFR was evaluated by the rate inulin-FITC clearance as previously described (91, 92). Inulin-FITC (Millipore-Sigma, Cat. No. F3272) was dissolved in 0.9% NaCl (5% w/v), and the solution was dialyzed with a 1,000-kDa dialysis membrane at room temperature in the dark for 24 h, followed by sterile filtering through a 0.22-µm filter. Mice were briefly anesthetized under isoflurane and FITC-inulin (2 µL/g body weight) was injected into the retroorbital sinus. Blood was then collected in heparinized capillary tubes via a ∼1-mm tail snip at 3, 5, 7, 10, 15, 35, 56, and 75 minutes following FITC-inulin injection. During blood collection, the mouse was conscious and allowed free movement within their cage. Blood was centrifuged for 10 minutes at 4,000 rpm at 4°C, the resulting plasma was collected and diluted (1:20) in 0.5 mol/L of HEPES buffer (pH 7.4) and loaded into a 96-well plate. The fluorescence was detected using a BioTek Synergy II plate reader against a FITC-inulin standard curve. GFR was calculated using a two-phase exponential decay curve fit in GraphPad Prism. At euthanasia, blood was collected via cardiac puncture, allowed to clot, and centrifuged at 4000xGfor 10 min at 4°C. The resulting serum was diluted (1:25 in DiH_2_O) and BUN was measured using a commercial assay kit (Arbor Assays, Cat. No. K024).

### Animal model of peripheral artery disease

Femoral artery ligation (FAL) (39, 94) was performed by anesthetizing mice with intraperitoneal injection of ketamine (90 mg/kg) and xylazine (10 mg/kg) and surgically inducing unilateral hindlimb ischemia by placing silk ligatures on the femoral artery just distal the inguinal ligament and immediately proximal to the saphenous and popliteal branches. Buprenorphine (0.05 mg/kg) was given post-operatively for analgesia.

### Limb perfusion assessment

Limb perfusion was assessed by laser Doppler flowmetry (moorVMS-LDF, Moor Instruments) prior to surgery, post-surgery, three days post-surgery, seven days post-surgery, and just prior to euthanasia as described previously (39, 69, 95). Both hindlimbs were shaved and the laser Doppler probe was placed ∼1-2mm away from the middle of the posterior side of the paw and the posterior side of the lateral head of the gastrocnemius muscle. Perfusion recovery was reported as a percentage of the non-ischemic limb.

### Assessment of Hindlimb Grip Strength

Unilateral hindlimb grip strength was measured using a Grip Strength Test Instrument (BIOSEB; Model No. BIO-GS3) prior to surgery, three days post-surgery, seven days post-surgery, and just prior to euthanasia. The mice were allowed to firmly grip a metal T-shaped bar with a single hindlimb paw and then were pulled straight back with increasing force until the mouse released the bar. Three trials were performed on both the control and surgical limb. The trial with the highest force was used for analysis and grip strength was reported as percentage of non-surgical control limb.

### Nerve-Mediated Muscle Contraction

The maximal twitch and tetanic force levels of the tibialis anterior (TA) muscle were measured *in-situ* by stimulation of the sciatic nerve as previously described (87, 96). Briefly, mice were anaesthetized with an intraperitoneal injection of xylazine (10 mg/kg) and ketamine (100 mg/kg). The distal tendon was tied using a 4-0 silk suture attached to the lever arm of the force transducer (Cambridge Technology; Model No. 2250). Muscle contractions were elicited by stimulating the sciatic nerve via bipolar electrodes using square wave pulses of 0.02ms (Aurora Scientific, Model 701A stimulator). Lab-View-based DMC program (version v5.500) was used for data collection and servomotor control. After optimal length of the muscle was obtained using twitch contractions, three isometric tetanic forces were performed using 500ms supramaximal electrical pulses at a stimulation frequency of 150Hz with at least one minute of rest between contractions. The highest force among the three measurements was reported as the peak tetanic force. Following tetanic contractions, three isometric twitch contractions (1Hz) were performed. Peak twitch and tetanic force levels were reported as absolute and specific (normalized to muscle weight) force levels.

### Isolation of Skeletal Muscle Mitochondria

Skeletal muscle mitochondria were isolated as previously described (38, 93). The gastrocnemius was dissected, cleaned of connective tissue, and minced into a fine paste on ice. Next, the minced muscle was incubated in ice cold mitochondrial isolation media (MIM) (50mM MOPS, 100mM KCl, 1mM EGTA, 5mM MgSO_4_) containing 0.025% w/v trypsin for five minutes, followed by centrifugation at 500xG for five minutes at 4°C. The resulting supernatant containing trypsin was decanted and the pellet was resuspended with MIM containing 0.02% w/v bovine serum albumin (BSA 2g/L). The sample was then homogenized on ice using a glass-Teflon homogenizer and subsequently centrifuged at 800xG for ten minutes at 4°C. The resulting supernatant was collected and centrifuged again at 10,000xG at 4°C resulting in a mitochondrial rich pellet. The pellet was washed with MIM to remove damaged mitochondria and gently resuspended in MIM without BSA. Protein concentration of the resuspension was assessed using bicinchoninic acid protein assay (ThermoFisher Scientific, Cat. No. A53225)

### Assessment of Mitochondrial Oxygen Consumption

High-resolution respirometry was performed using an Oroboros Oxygraph-2k (O2K) to measure oxygen consumption (*J*O2) at 37°C. Twenty micrograms of mitochondria were added to the O2K chamber in 2ml of buffer D (105 mM K-MES, 30 mM KCl, 1 mM EGTA, 10 mM K_2_HPO_4_, 5 mM MgCl_2_-6H_2_O, 2.5 mg/mL BSA, pH 7.2) supplemented with 5 mM creatine monohydrate. Mitochondria were energized by the addition of 5mM pyruvate, 2.5mM malate, and 0.2mM octanoylcarnitine. Next we added a clamp system containing ATP (5mM), phosphocreatine (PCr) (1mM), and creatine kinase (CK) (20 U/ml) which couples the interconversion of ATP and ADP to that of phosphocreatine (PCr) and free creatinine, to titrate the extra mitochondrial ATP/ADP ratio, thus free energy of ATP hydrolysis (ΔG_ATP_), to measure mitochondrial oxygen consumption at physiologically relevant levels of energy demand as done previously (97). Exogenous cytochrome c (10mM) was used to assess outer-membrane integrity of isolated mitochondria and samples with more than a 25% increase in oxygen consumption were excluded from this study. The ΔG_ATP_ was plotted against the corresponding *J*O2, and the slope was used to represent conductance throughout mitochondrial oxidative phosphorylation (OXPHOS), where lower conductance indicates impaired mitochondrial energetics.

### Skeletal Muscle Morphology and Ischemic Lesion Area

The tibialis anterior muscle (TA) from both limbs was removed, embedded in optimal cutting temperature (OCT) compound, and immediately frozen in liquid nitrogen-cooled isopentane for cryosectioning. Using a Leica 3050S cryotome, 10µm-thick transverse sections of the TA were cut and mounted on microscope slides. Skeletal muscle morphology and ischemic lesion area was assessed using light microscopy and standard methods of hematoxylin and eosin staining (H&E). Slides were imaged at x20 magnification with an Evos FL2 Auto microscope (ThermoFisher Scientific), and tiled images of the entire section were obtained for analysis in using Image J software. The ischemic lesion area was quantified by manually tracing areas of the muscle section with signs of ischemic injury (necrotic or regenerating myofibers). The non-myofiber area was measured by thresholding the image to quantify the area tissues between myofibers which was expressed as percentage of the total section area. Regenerating myofibers were quantified by manually counting myofibers that contained centralized nuclei by a blinded investigator.

### Perfused and Total Muscle Capillary Density

Mice received a retro-orbital injection of 50 µL of 1 mg/mL Griffonia simplicifolia lectin (GSL) isolectin B4, Dylight 649 (Vector Laboratories; Cat. No. DL-1208) to fluorescently label α-galactose residues on the surface of endothelial cells of perfused capillaries. Following the injection, animals were returned to their cage and allowed 1-2 hours of free movement prior to euthanasia and muscle harvest. The tibialis anterior muscle was frozen in OCT compound and cryo-sectioned as described above. Transverse muscle sections were fixed with 4% paraformaldehyde (ThermoFisher Scientific, Cat. No. J19943-K2) for ten minutes, and permeabilized with 0.25% triton X-100 (Millipore-Sigma, Cat. No. 93443). Following three washes with PBS, slides were blocked in PBS + 5% goat serum + 1% BSA for 4- 6 hours. Total capillaries were labeled with a primary antibody raised against PECAM1 (anti- CD31, Abcam, Cat. No. ab28364, 1:100 dilution in blocking solution) overnight at 4°C. The following day slides were washed with PBS and incubated for one hour at room temperature with Alexa-Fluor555 anti-rabbit secondary antibody (ThermoFisher Scientific, Cat. No. A11034, 1:250 dilution) and wheat germ agglutin (WGA) conjugated with Alexa-Flour488 1mg/ml (ThermoFisher Scientific, Cat. No. W11261, 1:100 dilution) to label myofiber membranes. Next, slides were washed with PBS and subsequently cover slipped with Vectashield Hardmount containing DAPI (Vector Laboratories, Cat. No. H-1500-10). Images were obtained at x20 magnification using an Evos FL2 Auto microscope (ThermoFisher Scientific) and tiled/merged images of the entire muscle section were used for analysis. The number of perfused and total capillaries density measurements were quantified on thresholded images using a particle counter in Fiji/ImageJ and the quantified results were expressed as percent perfused and total number per area of the muscle section. Skeletal myofiber cross-sectional area (CSA) was determined using MuscleJ (7), an automated analysis software developed in Fiji.

### Primary Endothelial Cell Culture

Primary Endothelial Cells were isolated from the liver of 4-week-old male and female C57BL6J mice (Jackson Laboratory, Stock #000664). All experiments were performed in three biologically independent samples. Primary cells were grown to ∼80% confluency on 0.25% gelatin (ScienCell, Cat. No. 0423) coated flasks in endothelial cell growth medium (PromoCell, Cat. No. C-22022) supplemented with 1% penicillin/streptomycin (ScienCell, Cat. No. 0503) using standard culture conditions (37°C with 5% CO_2_). Cells were stained with anti-CD31 antibody (Abcam, Cat. No. ab28364, 1:100 dilution in blocking solution) and DAPI to validate that the cells were CD31-positive. Hypoxia and nutrient deprivation was induced by replacing culture media with Hank’s balanced salt solution (HBSS; ThermoFisher Scientific; Cat. No. 24020) and placing cells within a cake pan hypoxia chamber flushed with nitrogen gas for ten minutes prior to sealing as previously described (98). To activate the AHR, cells were treated with indoxyl sulfate (100μM, Millipore-Sigma, Cat. No. 13875) or vehicle control DMSO (ThermoFisher Scientific; Cat. No. BP231).

### RNA isolation and quantitative PCR.

Total RNA was isolated from primary murine endothelial cells and mouse extensor digitorum longus muscle using TRIzol (Invitrogen, Cat. No. 15-596-018) and a Direct-zol RNA MiniPrep kit (Zymo Research, Cat. No. R2052). RNA concentration and purity were assessed using spectrophotometery. cDNA was generated from 500ng of RNA (tissue samples) or 100ng RNA (cell culture experiments) using the LunaScript RT Supermix kit (New England Biolabs, Cat. No. E3010L) according to the manufacturer’s directions. Real-time PCR (RT-PCR) was performed on a Quantstudio 3 (ThermoFisher Scientific) using Luna Universal qPCR master mix for Sybr Green primers (New England Biolabs, Cat. No. M3003X). All primer sequences are listed in Supplementary Table 1. Relative mRNA expression was calculated using 2^-ΔΔCT^ from the relevant control group.

### RNA sequencing.

Library preparation and RNA sequencing was performed by Quick Biology, Inc. (Pasadena, CA). RNA integrity was checked by Agilent Bioanalyzer 2100; only samples with clean rRNA peaks were used. Library for RNA-Seq was prepared according to KAPA Stranded mRNA-Seq poly(A) selected kit with 201-300bp insert size (KAPA Biosystems, Wilmington, MA) using 250 ng total RNA as input. Final library quality and quantity was analyzed by Agilent Bioanalyzer 2100 and Life Technologies Qubit3.0 Fluorometer. 150 bp paired end reads were sequenced on Illumina HighSeq 4000 (Illumina Inc., San Diego, CA). The reads were first mapped to the latest UCSC transcript set using Bowtie2 (99) and the gene expression level was estimated using RSEM (100) TMM (trimmed mean of M-values) was used to normalize the gene expression. Differentially expressed genes were identified using the edgeR program (101). Genes showing altered expression with *P* < 0.05 and more than 1.5-fold changes were considered differentially expressed. Goseq was used to perform the gene ontology (GO) enrichment analysis.

### Study Approval

All procedures were approved by the Institutional Animal Care and Use Committee of the University of Florida (Protocol 201810484).

### Data Availability

All source data are available from the corresponding author upon request. RNA sequencing data have been deposited to the Gene Expression Omnibus (https://www.ncbi.nlm.nih.gov/geo/) under accession number GSE234508.

### Statistical Analysis

All data are presented as the mean ± standard deviation (SD). Normality of data was tested with the Shapiro-Wilk test and/or inspection of QQ plots. Data involving comparisons of two groups were analyzed using a student’s t-test when normally distributed and Mann-Whitney test when normality could not be assessed. When making comparisons with more than two groups, data were analyzed using two-way ANOVA with Sidak’s post hoc testing for multiple comparisons when significant interactions were detected. Mixed-effects analysis and three-way ANOVA were used to determine differences when three or more main effects were present. In all cases, *P* < 0.05 was considered statistically significant. All statistical testing, except for RNA sequencing analysis, was conducted using GraphPad Prism software (version 9.0).

## Acknowledgements

This study was supported by National Institutes of Health (NIH) grant R01-HL149704 (T.E.R.). The content is solely the responsibility of the authors and does not necessarily represent the official views of the National Institutes of Health.

## Author Contributions

VRP and TER conceived the study and designed the experiments. VRP, JT, QY, JM, OL, and TER performed experiments and collected data. VRP, JT, QY, JM, OL, and TER analyzed and interpreted the data. VRP and TER drafted the manuscript. All authors edited, revised, and approved the final version of the manuscript.

