## Supplemental Material for "Activation of the Aryl Hydrocarbon Receptor in Endothelial Cells Impairs Ischemic Angiogenesis in Chronic Kidney Disease"

**Running Head:** AHR Activation Impairs Ischemic Angiogenesis

**Supplemental Table 1. Primers for qPCR and DNA recombination**

| Species | Gene | Chemistry | Sequence |
|---|---|---|---|
| Mus Musculus | *Ahr* | Sybr Green | 5'– TTCCAGGTTCTCAGGCATTC – 3'<br>5'– TGGGAGCTACAGGAATCCAC – 3' |
| Mus Musculus | *Cyp1a1* | Sybr Green | 5'– CAGCCTTCCCAAATGGTTTA – 3'<br>5'– GCCTGGGCTACACAAGACTC – 3' |
| Mus Musculus | *Ahrr* | Sybr Green | 5'– GACTTCTGCAGACAGCTACA – 3'<br>5'– TGTCAAGAAGGCCGAGTACT – 3' |
| Mus Musculus | *Vegfa* | Sybr Green | 5' – CGAGATAGAGTACATCTTCAAGCC – 3'<br>5' – TCATCGTTACAGCAGCCTGC – 3' |
| Mus Musculus | *Vegf121* | Sybr Green | 5' – TGCAGGCTGCTGTAACGATG – 3'<br>5' – CCTCGGCTTGTCACATTTTTCT – 3' |
| Mus Musculus | *Vegf165* | Sybr Green | 5' – TGCAGGCTGCTGTAACGATG – 3'<br>5' – GAACAAGGCTCACAGTGATTTTCT – 3' |
| Mus Musculus | *Angpt1* | Sybr Green | 5' – CATTCTTCGCTGCCATTCTG – 3'<br>5' – GCACATTGCCCATGTTGAATC – 3' |
| Mus Musculus | *Hif1a* | Sybr Green | 5' – GGGGAGGACGATGAACATCAA – 3'<br>5' – GGGTGGTTTCTTGTACCCACA – 3' |
| Mus Musculus | *End1* | Sybr Green | 5' – GCACCGGAGCTGAGAATGG – 3'<br>5' – GTGGCAGAAGTAGACACACTC – 3' |
| Mus Musculus | *Nos2* | Sybr Green | 5' – CAGCTGGGCTGTACAAACCTT – 3'<br>5' – CATTGGAAGTGAAGCGTTTCG – 3' |
| Mus Musculus | *Egf* | Sybr Green | 5' – AGCATCTCTCGGATTGACCCA – 3'<br>5' – CCTGTCCCGTTAAGGAAAACTCT – 3' |
| Mus Musculus | *Cox7a1* | Sybr Green | 5' – GCTCTGGTCCGGTCTTTTAGC – 3'<br>5' – GTACTGGGAGGTCATTGTCGG – 3' |
| Mus Musculus | *Atp5k* | Sybr Green | 5' – GTTCAGGTCTCTCCACTCATCA – 3'<br>5' – CGGGGTTTTAGGTAACTGTAGC – 3' |
| Mus Musculus | *Atp5d* | Sybr Green | 5' – TGCTTCAGGCGCGTACATAC – 3'<br>5' – CACTTGCTTGACGTTGGCA – 3' |
| Mus Musculus | *Tfam* | Sybr Green | 5' – ATTCCGAAGTGTTTTTCCAGCA – 3'<br>5' – TCTGAAAGTTTTGCATCTGGGT – 3' |
| Mus Musculus | *Sod2* | Sybr Green | 5' – ACAAACCTGAGCCCTAAGGGT – 3'<br>5' – GAACCTTGGACTCCCACAGAC – 3' |
| Mus Musculus | *Actb* | Sybr Green | 5' – GGCTGTATTCCCCTCCATCG– 3'<br>5' – CCAGTTGGTAACAATGCCATGT– 3' |
| Mus Musculus | *L32* | Sybr Green | 5'– TTCCTGGTCCACAATGTCAA – 3'<br>5'– GGCTTTTCGGTTCTTAGAGGA – 3' |

**Primers used for DNA recombination**

| Species | Gene | Sequence |
|---|---|---|
| Mus Musculus | *Ahr* | 5'– ATCTTGTGTCAGGAACAGGCCATC – 3'<br>5'– GGTACAAGTGCACATGCCTGC – 3' |

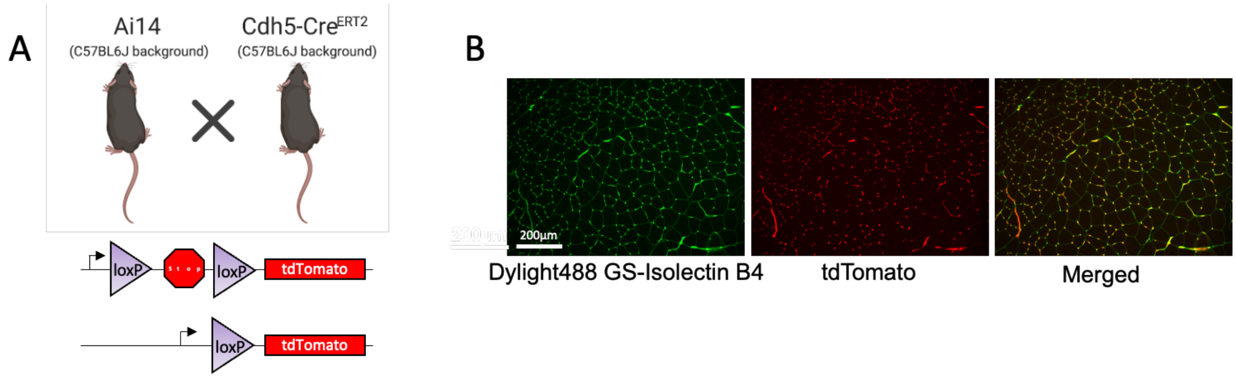


**Supplemental Figure 1. Validation of Endothelial-specific Cre expression via reporter mouse.** (A) The tamoxifen inducible Cdh5-Cre$^{ERT2}$ (Taconic) mice were and bred with Ai14 (Jackson Laboratory) mice for validation of Cre specificity. (B) Offspring injected with tamoxifen showed robust and specific expression of the TdTomato fluorescent protein in the capillaries of the tibialis anterior muscle.

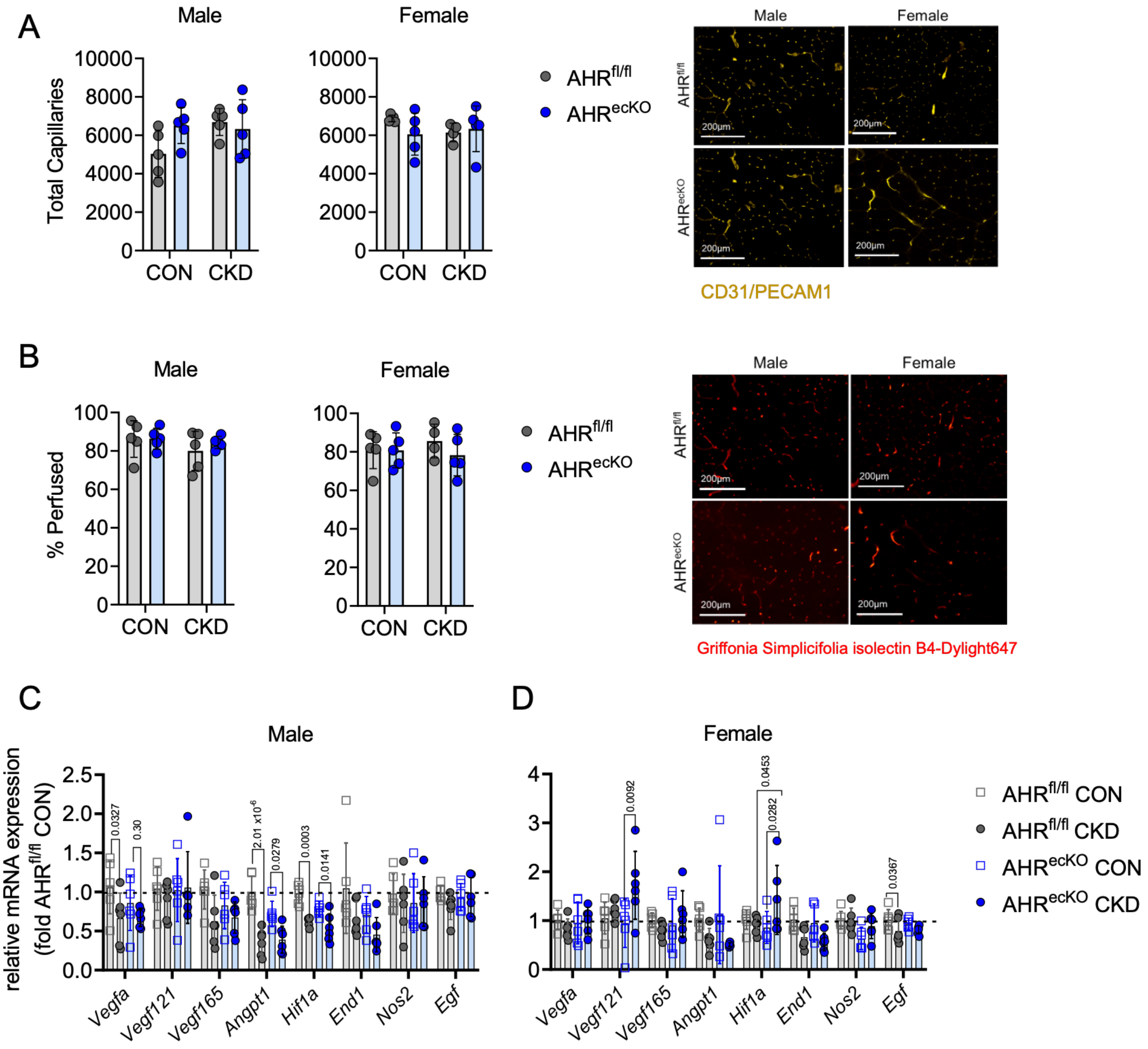


**Supplemental Figure 2. Endothelium-specific Deletion of the AHR Does not Alter Capillary Densities in the Non-ischemic Limb.** (A) Total number of capillaries and (B) perfused capillaries measured by retroorbital injection of isolectin and quantified as a percentage of total capillaries. All capillary density measurements were performed on a cross section of tibialis anterior muscle of the non-ischemic limb (n=5/group/sex). (C) expression of angiogenic and vasoreactive genes in non-ischemic extensor digitorum longus muscle from male and (D) female mice (n=6/group/sex). Statistical analyses were performed in panels A and B using Mann-Whitney test (normality could not be established) and panels C and D using two-way ANOVA with Sidak's post hoc testing for multiple comparisons. Error bars represent the standard deviation.

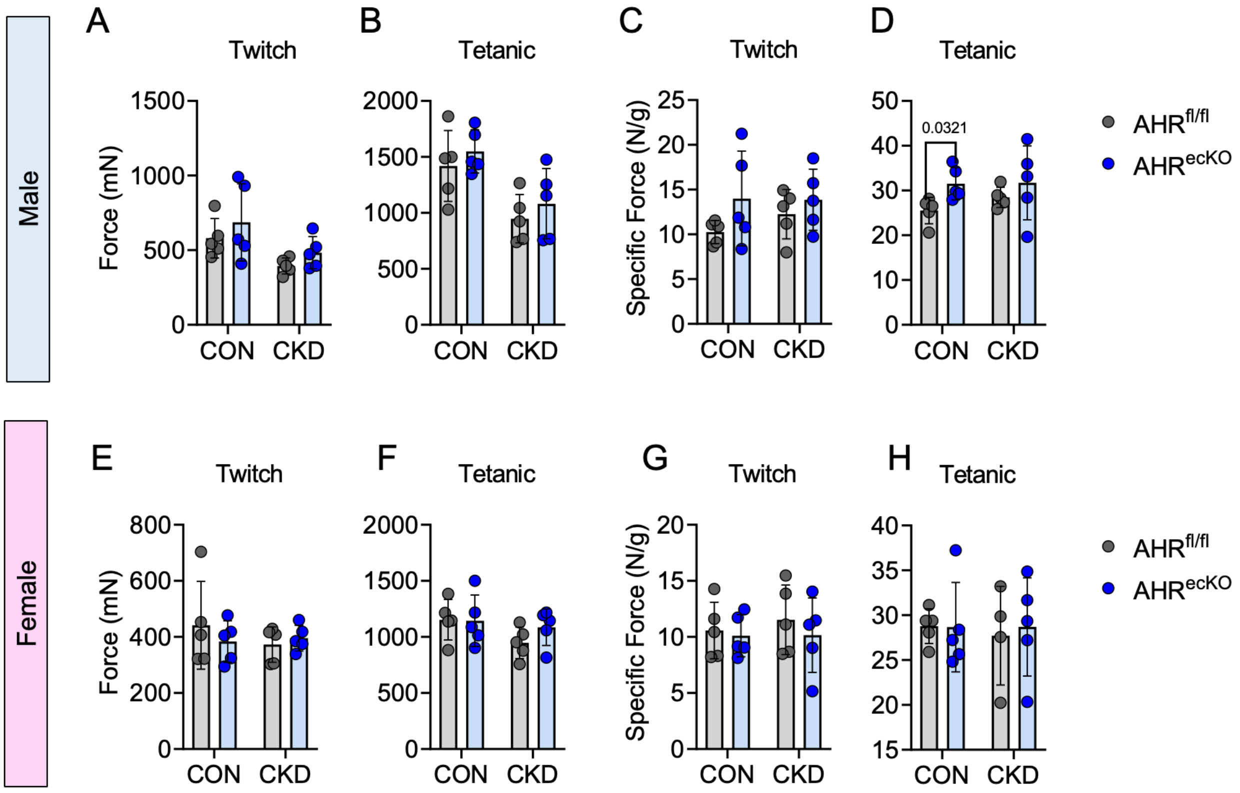


**Supplemental Figure 3. Endothelium-Specific AHR Deletion Does Not Improve Non-Ischemic Muscle Function in Mice with CKD.** (A-H) Muscle contractile function quantification generated using nerve mediated *in-situ* functional analyses of the non-ischemic tibialis anterior muscle (n=5/group/sex). (A) Maximal absolute force generated from a twitch contraction (1Hz stimulus) and (B) tetanic contraction (150Hz stimulus) in male mice. (C) Muscle contractile force from maximal twitch and (D) tetanic contractions normalized to muscle wet weight in males. (E) Maximal absolute force generated from a twitch contraction (1Hz stimulus) and (F) tetanic contraction (150Hz stimulus) in female mice. (G) Muscle contractile force from maximal twitch and (H) tetanic contractions normalized to muscle wet weight in female mice. Data were analyzed using a Mann-Whitney test (normality could not be established). Error bars represent the standard deviation.

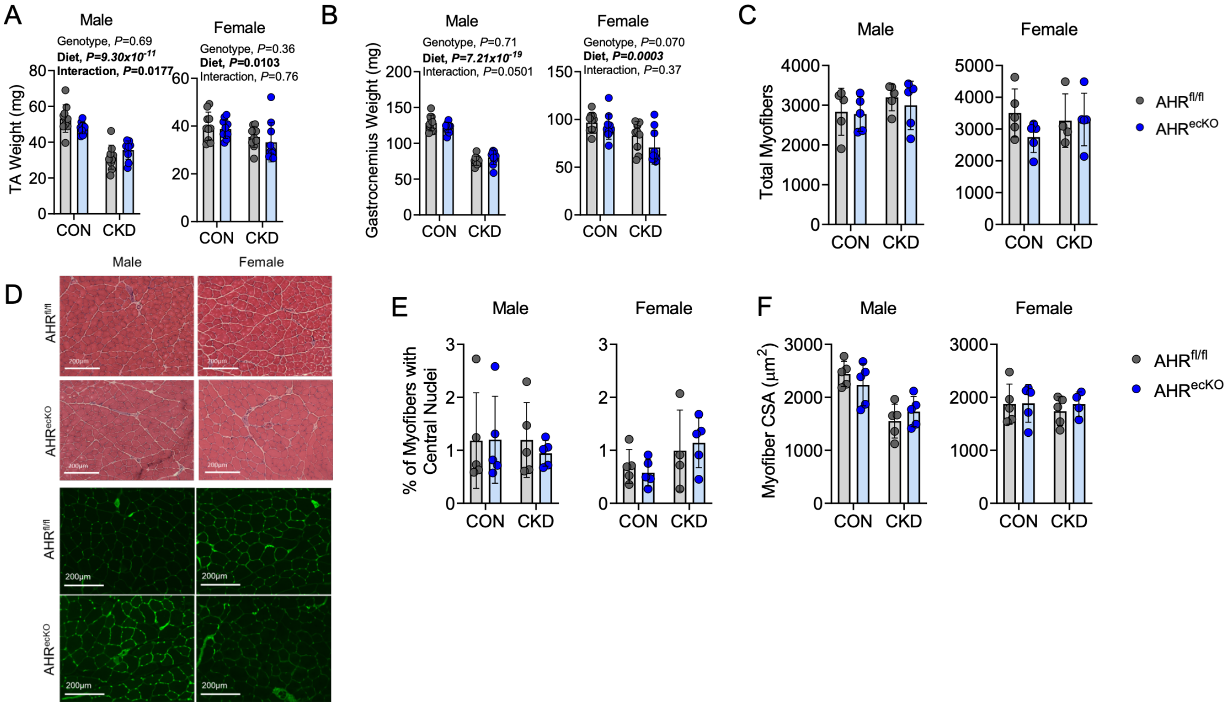


**Supplemental Figure 4. Endothelium-Specific AHR Deletion Has No Impact on Muscle Size or Histopathology in Non-Ischemic Muscle.** (A) Weights of the tibialis anterior and (B) gastrocnemius muscles (n=10/group/sex). (C) Total number of fibers in the tibialis anterior of non-ischemic limb (n=5/group/sex). (D) Representative 20x images of hematoxylin & eosin staining and laminin staining immunofluorescence of cross sections of the non-ischemic tibialis anterior muscle. (E) Percentage of total myofibers with centralized nuclei and (F) mean myofiber cross-sectional area of the non-ischemic tibialis anterior (n=5/group/sex). Panels A and B were analyzed using two-way ANOVA with Sidak's post hoc testing for multiple comparisons. Panels C-F were analyzed using a Mann-Whitney test (normality could not be established). Error bars represent the standard deviation.

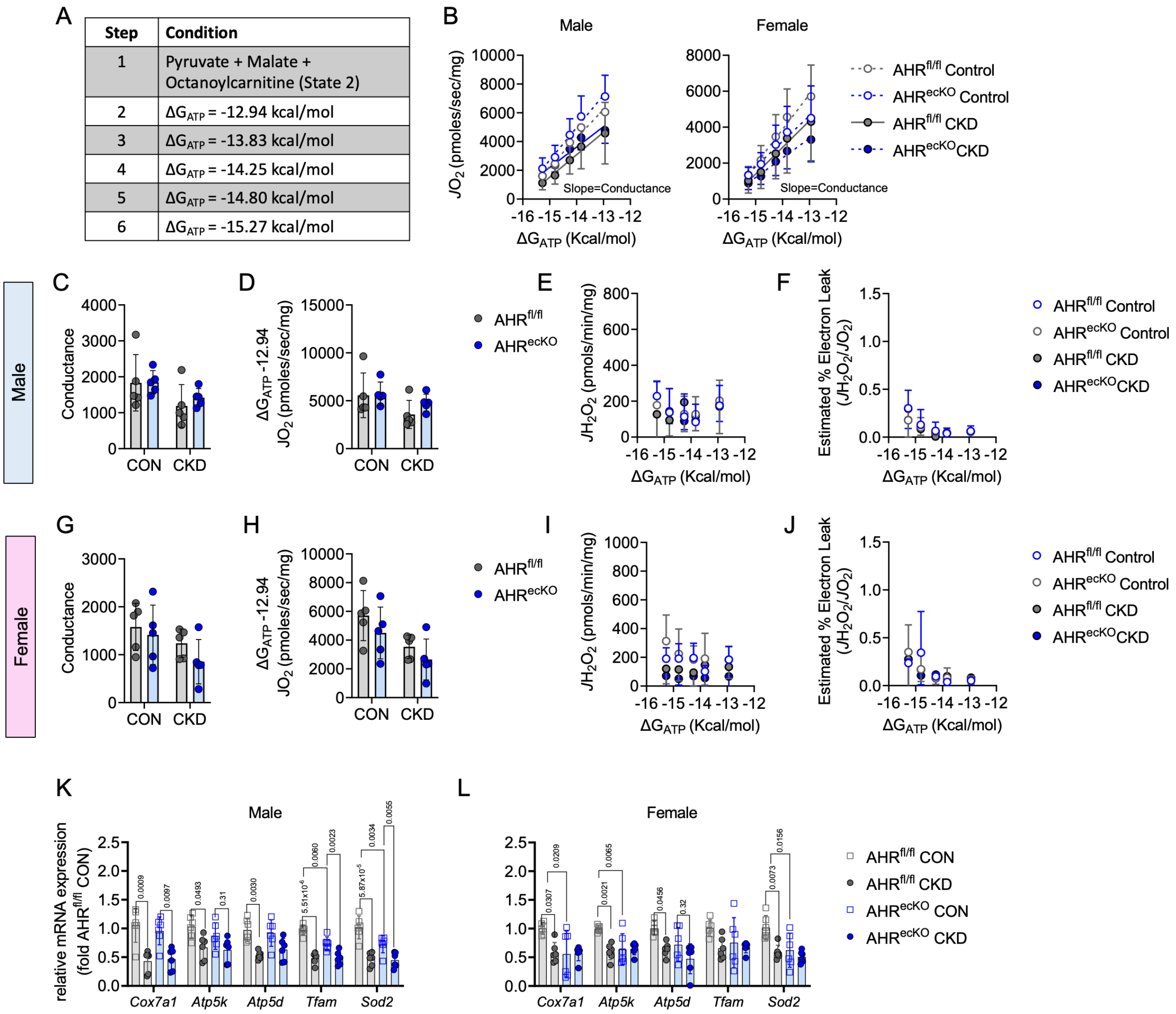


**Supplemental Figure 5. Endothelium-Specific AHR Deletion Does Not Impact Mitochondrial Function in Non-Ischemic Muscle.** Mitochondrial function in the non-ischemic gastrocnemius muscle (n=5/group/sex) was assessed using (A) a creatine kinase clamp system to measure oxygen consumption ($JO_2$) at physiologically relevant energy demands ($\Delta G_{ATP}$) fueled with pyruvate, malate and octanoylcarnitine. (B) Relationship between $JO_2$ and $\Delta G_{ATP}$ (C) quantification of the conductance (slope of $JO_2$ and $\Delta G_{ATP}$ relationship) and (D) $JO_2$ at the highest energy demand ($\Delta G_{ATP}$ -12.94) in male mice. (E) Mitochondrial $H_2O_2$ production at each $\Delta G_{ATP}$ and (F) Estimated percent electron leak ($JH_2O_2$/ $JO_2$) at each $\Delta G_{ATP}$ in males. (G) quantification of the conductance (slope of $JO_2$ and $\Delta G_{ATP}$ relationship) and (H) $JO_2$ at the highest energy demand ($\Delta G_{ATP}$ -12.94) in female mice. (I) Mitochondrial $H_2O_2$ production at each $\Delta G_{ATP}$ and (J) Estimated percent electron leak ($JH_2O_2$/ $JO_2$) at each $\Delta G_{ATP}$ in females. (K) expression of mitochondrial related genes in extensor digitorum longus muscle in male and (L) female mice (n=6/group/sex). Panels B-J were analyzed using a Mann-Whitney test (normality could not be established). Panels K and L were analyzed using two-way ANOVA with Sidak's post hoc testing for multiple comparisons. Error bars represent the standard deviation.
